# Transient intestinal colonization by a live-attenuated oral cholera vaccine induces protective immune responses in streptomycin-treated mice

**DOI:** 10.1101/2020.05.19.104471

**Authors:** Bolutife Fakoya, Brandon Sit, Matthew K. Waldor

## Abstract

Current mouse models for evaluating the efficacy of live oral cholera vaccines (OCVs) have important limitations. Conventionally raised adult mice are resistant to intestinal colonization by *Vibrio cholerae,* but germ-free mice can be colonized and have been used to study OCV immunogenicity. However, germ free animals have impaired immune systems and intestinal physiology; also, live OCVs colonize germ free mice for many months, which does not mimic the clearance kinetics of live OCVs in humans. Here, we leverage antibiotic-treated, conventionally raised adult mice to study the effects of transient intestinal colonization by a live OCV *V. cholerae* strain. In a single dose vaccination regimen, we found that HaitiV, a live-attenuated OCV candidate, was cleared by streptomycin treated adult mice within a week after oral inoculation. This transient colonization elicited far stronger adaptive immune correlates of protection against cholera than did inactivated whole-cell HaitiV. Infant mice from HaitiV vaccinated dams were also significantly protected from choleric disease than pups from inactivated-HaitiV dams. Our findings establish the benefits of antibiotic treated mice for live OCV studies as well as its limitations and underscore the immunogenicity of HaitiV.

**Importance:** Oral cholera vaccines (OCVs) are being deployed to combat cholera but current killed OCVs require multiple doses and show little efficacy in young children. Live OCVs have the potential to overcome these limitations but small animal models for testing OCVs have shortcomings. We used an antibiotic treatment protocol for conventional adult mice to study the effects of short-term colonization by a single dose of HaitiV, a live OCV candidate. Vaccinated mice developed vibriocidal antibodies against *V. cholerae* and delivered pups that were resistant to cholera, whereas mice vaccinated with inactivated HaitiV did not. These findings demonstrate HaitiV’s immunogenicity and suggest that this antibiotic treatment protocol will be useful for evaluating the efficacy of live OCVs.

## Introduction

*Vibrio cholerae* is the cause of cholera and, following ingestion of water or food contaminated with this Gram-negative rod, humans can develop the severe and sometimes lethal dehydrating diarrhea that characterizes cholera. Cholera remains a major threat to global public health, with approximately 2.9 million cases and 95,000 deaths reported annually (1). Serologic classification of *V. cholerae* is based on the structure and chemistry of the abundant LPS O-antigen and the O1 serogroup of *V. cholerae* has given rise to all cholera pandemics. The O1 serogroup is further subdivided into Ogawa and Inaba serotypes that differ by the presence or absence of methylation of the terminal perosamine on their respective O-antigens (2). Current pandemic cholera is predominantly caused by an O1 ‘variant’ El Tor biotype strain, such as the strain responsible for the Haitian epidemic in 2010 (3).

Prior exposure to *V. cholerae* can elicit long-lived protective O-antigen-specific responses against *V. cholerae* (4, 5), suggesting that vaccination has the potential to elicit protective immunity if vaccines can safely mimic elements of natural infection. As such, several vaccination strategies for cholera have been developed over the decades. Among these, oral cholera vaccines (OCV) are attractive options as they stimulate immunity at the intestinal mucosal surface, the site of infection, and because of their ease of administration. Killed whole-cell OCVs, such as Shancol, are being increasingly adopted as frontline public health tools both in endemic regions (6) as well as to limit outbreaks (7).Live-attenuated OCVs have also been developed and have theoretical advantages over killed OCVs including *in vivo* replication which enables continuous presentation of *in vivo*-induced antigens at the intestinal mucosal surface (8). In contrast to killed OCVs, live vaccines will likely offer single dose efficacy, a particularly important feature for reactive vaccination campaigns during epidemics. Furthermore, live OCVs appear to be more effective in children less than 5 years of age (9, 10), a group that is highly susceptible to death from cholera and that is not adequately protected by killed OCVs. In volunteer studies, these vaccines have shown great promise (11, 12), but none are approved for use in cholera endemic regions.

The development of both inactivated and live OCVs has been hampered by the lack of a small animal model that closely recapitulates cholera pathogenesis in the setting of normal immune reactivity. *V. cholerae* readily colonizes the intestines of infant mice and infant rabbits, reviewed in (13), where cholera-like disease can be observed, but these models lack mature immune systems. Conventionally raised adult mice cannot be orally colonized by *V. cholerae,* likely due to their resident gut microbiota (14). Germ free (GF) adult mice, which lack a microbiota, can be colonized by *V. cholerae* and have been used to profile OCV immunogenicity, but immune and intestinal physiological development is aberrant in these animals (15). Nonetheless, vaccinated GF mice develop immune correlates of clinical protection against toxigenic *V. cholerae,* including circulating vibriocidal antibodies and antigen specific antibody responses (14, 16, 17). Another limitation of the GF adult mouse model is that following oral administration of a live OCV, they remain consistently colonized by *V. cholerae* for periods exceeding 3 months (14, 17), making single-dose live OCV regimens difficult to interpret; moreover, the prolonged colonization in this model precludes challenging vaccinated animals with virulent *V. cholerae*. The consequence and significance of long-term monocolonization with a live OCV in GF mice also remains unknown.

Many studies have shown that oral administration of broad-spectrum antibiotics to mice depletes the gut microbiota and enables intestinal colonization by diverse bacteria (18). This is beneficial as antibiotic treated mice are conventionally raised and do not display the immunological and developmental defects that characterize GF mice. Antibiotic treatments can enable wild-type *V. cholerae* intestinal colonization for similar durations, typically 5-7 days, that humans are colonized by live OCVs (19, 20). We sought to leverage this model to profile HaitiV, a live-attenuated OCV derived from HaitiWT, a virulent *V. cholerae* O1 Ogawa clinical strain isolated during the Haitian cholera outbreak (21). HaitiV is non-toxigenic and highly engineered for biosafety and in infant rabbits HaitiV provides unprecedented rapid protection against virulent *V. cholerae* within 24 hours of administration (21); furthermore, we showed that this vaccine is immunogenic in GF mice (17).

Here, we adopted a streptomycin treated adult mouse model of *V. cholerae* (22) to profile HaitiV’s immunogenicity. Furthermore, we modified this model to assess the protective efficacy of immune responses in immunized female mice, by challenging their pups with HaitiWT. Our findings demonstrate the ease and utility of this approach for studies of live-attenuated OCVs.

## Results

### Vaccine inoculation protocol and colonization kinetics

We modified the protocol presented in Bueno et al (22) and orally treated 4-week-old C57BL/6 female mice with streptomycin (Sm) to deplete their intestinal microbiota to enable a longitudinal study of HaitV intestinal colonization and immunogenicity (Fig. 1A). Three days after initiating Sm treatment, mice were orally gavaged with either a single dose of 10^9^ CFU of HaitiV, an equivalent dose of formalin-inactivated HaitiV (FI-HaitiV) to mimic oral vaccination with an inactivated OCV, or sodium bicarbonate as a vehicle control. There were no apparent untoward effects of any of these regimens and over the course of the experiment, all mice gained weight (Fig. 1B).

**Figure 1.**
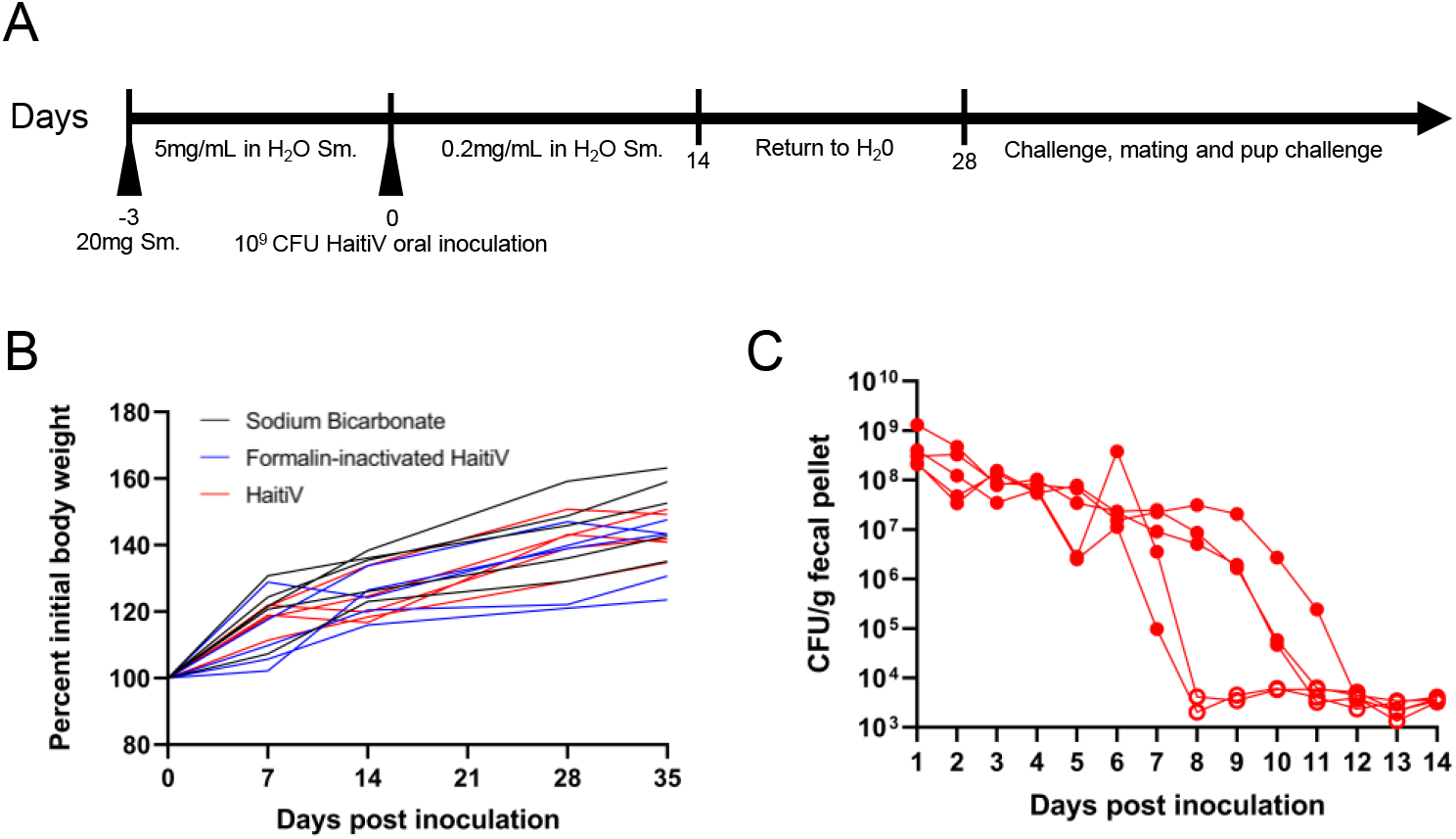
Transient intestinal colonization of HaitiV in Sm-treated adult mice following treatment with streptomycin. A) Schematic of streptomycin treatment and single oral inoculation with HaitiV. Black arrowheads indicate peroral treatment with either streptomycin in sodium bicarbonate, or oral gavage with bacteria. B) Bodyweight of all mice over the course of this study. C) Fecal shedding of HaitiV from mice inoculated with HaitiV; open symbols depict CFU levels below the limit of detection.

Plating of fresh fecal pellets (FP) from all animals inoculated with HaitiV revealed ~10^8^ CFU/g FP for 5-7 days post inoculation (dpi), suggesting that initially the vaccine robustly colonized the intestines of Sm treated adult mice. However, HaitiV was no longer detectable in FPs by 8-12 dpi (Fig. 1C, Fig. S1B), indicating clearance of the vaccine strain. No CFUs of HaitiV were recovered from Sm treated mice that had been orally inoculated with FI-HaitiV or buffer control. The clearance kinetics of HaitiV from these mice resembles that previously charted by Nygren et al (19) for wild type cholera toxin-producing *V. cholerae* isolates, suggesting that cholera toxin does not play a substantial role in intestinal colonization in this model.

### Transient HaitiV colonization elicits antibodies targeting multiple *V. cholerae* serotypes

Sera from all mice were individually assayed to quantify the circulating vibriocidal antibody titers (VATs) targeting Ogawa and Inaba *V. cholerae* strains. By 7dpi most mice did not have detectable VATs but by 14dpi when HaitiV was no longer detectable in FPs, most (3/5 mice) seroconverted and developed high circulating VATs against both serotyped matched (Ogawa) and serotype mismatched (Inaba) *V. cholerae* (Fig. 2A, Fig. S1C, Fig. S1D), though highest geometric mean VATs were directed against serotype matched (Ogawa) isolates (Fig. 2A, Fig. S1C). Only one mouse in the FI-HaitiV group developed circulating VATs against Ogawa serotype *V. cholerae,* and no mice in this group developed vibriocidal antibodies against Inaba *V. cholerae* in this group (Fig. 2B). None of the Sm treated mice that received sodium bicarbonate developed detectable vibriocidal antibodies. Thus, transient colonization by HaitiV is sufficient to elicit the generation of *V. cholerae* specific circulating markers of immunity.

**Figure 2.**
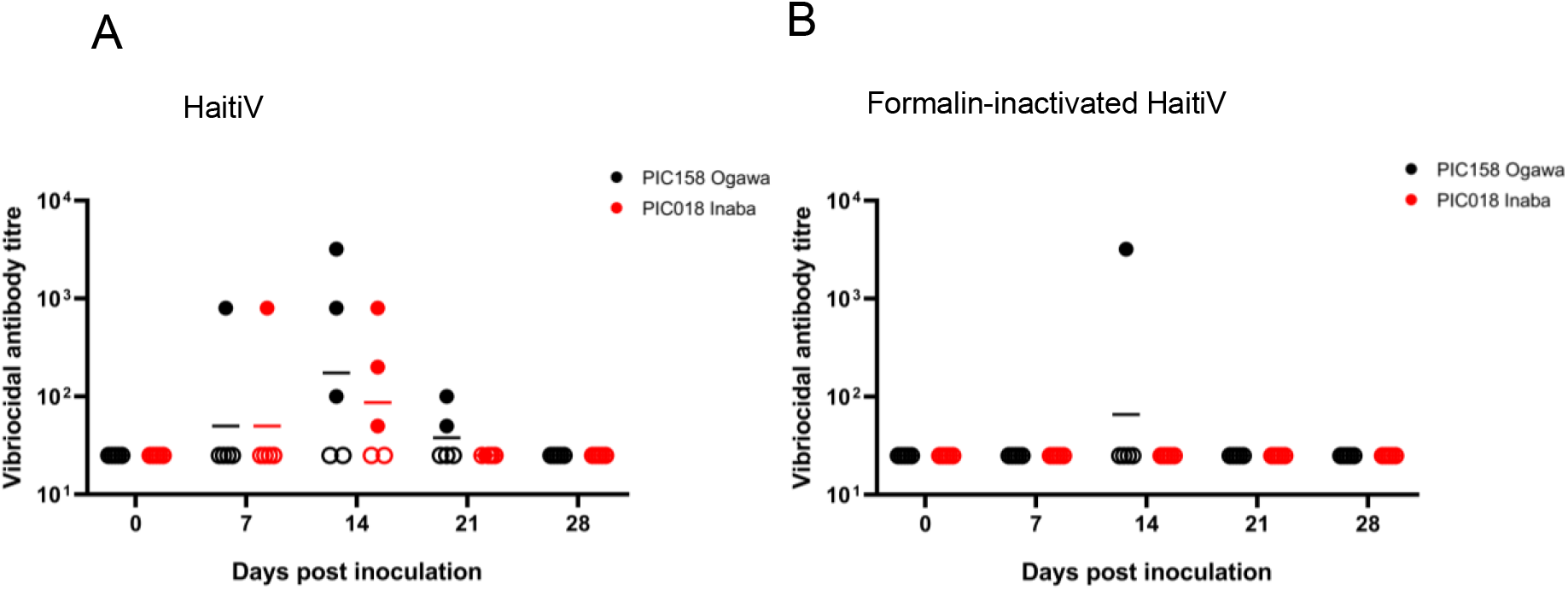
Serum vibriocidal antibody titers in Sm-treated mice immunized with HaitiV. Circles indicate the lowest dilutions at which specific vibriocidal activity was detected, and the height of the bars represent geometric mean titers in each group. Ogawa *V. cholerae* PIC158 was used to measure Ogawa serotype-specific antibodies (black) and Inaba *V. cholerae* PIC018 was used to measure Inaba serotype-specific antibodies (red). Titers below the limit of detection are indicated by open symbols.

### Streptomycin treated adult mice resist recolonization by *V. cholerae*

Administration of live-OCVs to GF mice leads to long term intestinal colonization (14, 17), precluding the possibility of challenge studies with wild-type *V. cholerae.* Since the Sm-treated mice cleared HaitiV, we investigated whether animals immunized with this OCV could be successfully challenged with HaitiWT. At 28dpi, when all mice had stopped shedding HaitiV, they (live HaitiV, FI-HaitiV, and vehicle control) were orally treated with sulfamethoxazole and trimethoprim (SXT) to reduce the intestinal microbiota (Fig. 3A); unlike HaitiV, HaitiWT is resistant to SXT. All three groups of mice from above and another group, specific pathogen free (SPF) mice that had not been Sm-treated or exposed to HaitiV was also included in this experiment, to test whether SXT treatments modify susceptibility to HaitiWT colonization. All 4 groups of mice were then orally gavaged with 10^9^ CFU of HaitiWT, and their FPs monitored for HaitiWT colonization. Unexpectedly, the three groups of mice (HaitiV, FI-HaitiV, and sodium bicarbonate) that had been previously treated with Sm failed to be colonized by HaitiWT (Fig. 3B). In contrast, the SXT-treated SPF mice were robustly colonized by HaitiWT (~10^8^ CFU/g FP) (Fig. 3B), indicating that although SXT treatment does susceptibilize mice to *V. cholerae* colonization, the prior Sm treatment of the vaccinated animals rendered them resistant to recolonization with *V. cholerae*.

**Figure 3.**
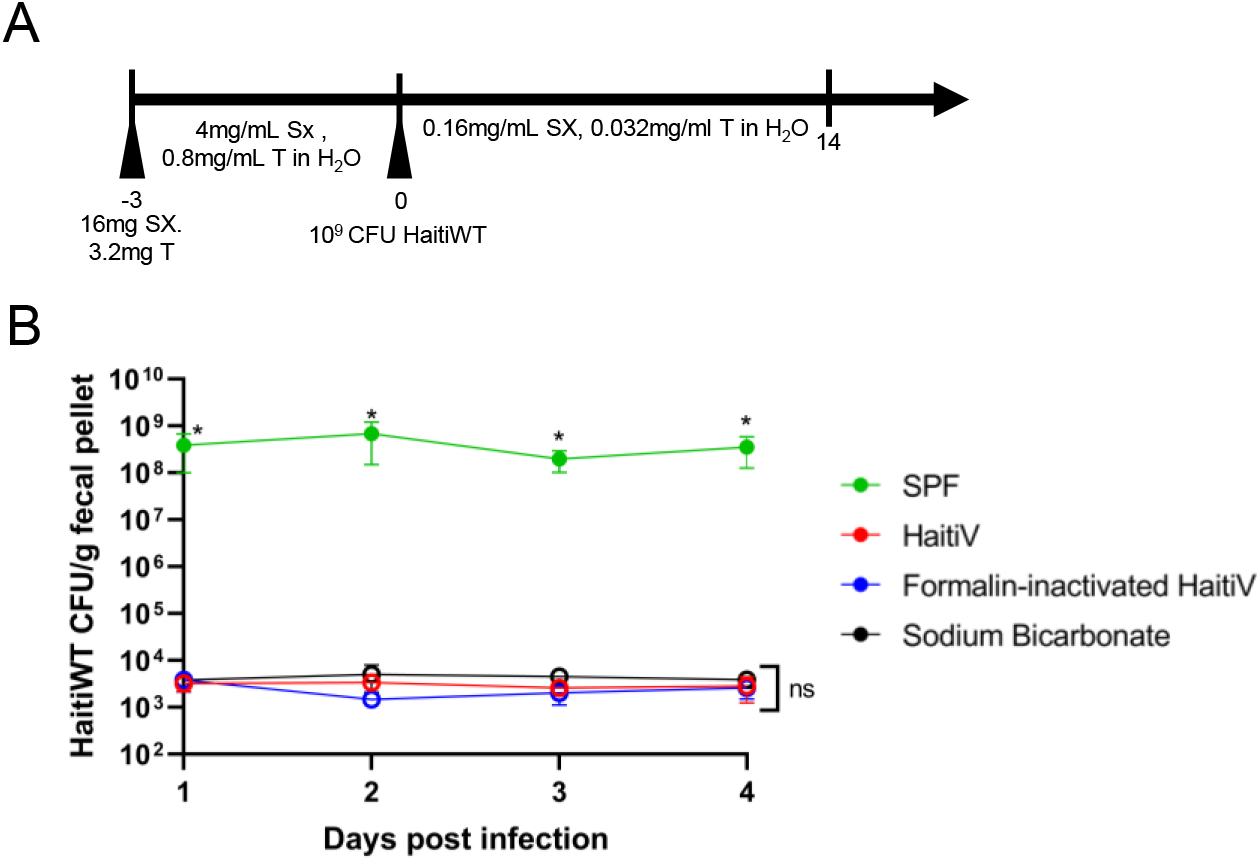
SXT treatment and re-colonization of Sm-treated adult mice by *V. cholerae.* A) All mice were treated with sulfamethoxazole and trimethoprim (SXT) and dosed with 10^9^ CFU of HaitiWT on day 0. B) Fecal shedding of HaitiWT from all mice; open symbols represent fecal pellets from which no CFUs of HaitiWT could be detected. An asterisk denotes differences with a P value of <0.05 as determined by a Mann Whitney U test.

Although the FPs of the previously Sm treated and HaitiWT challenged mice lacked detectable HaitiWT on the Sm agar plates used to detect *V. cholerae,* these plates contained high numbers of colonies of non-*V. cholerae* bacteria. 16s rRNA sequencing of Sm resistant small colonies showed that these colonies corresponded to either *Escherichia coli* or SXT resistant *Lactobacillus murinus,* which together were present at ~10^8^ CFU/g FP. These observations suggest that a bloom of Sm-resistant organisms and other changes in the gut microbiota associated with prior oral Sm treatment render mice resistant to *V. cholerae* colonization.

### Offspring of vaccinated dams are protected from virulent *V. cholerae*

As challenge studies could not be performed in Sm treated adult mice, we turned to the suckling mouse model of cholera to assess the protective efficacy of the immune response that was elicited by the transient HaitiV intestinal colonization. We recently found that the survival of suckling mice in this lethal challenge model can be used to gauge OCV efficacy (17). As such, animals from HaitiV, FI-HaitiV, and sodium bicarbonate groups were mated with SPF males and their neonatal pups infected with a lethal dose of HaitiWT (Fig. 4A).

**Figure 4.**
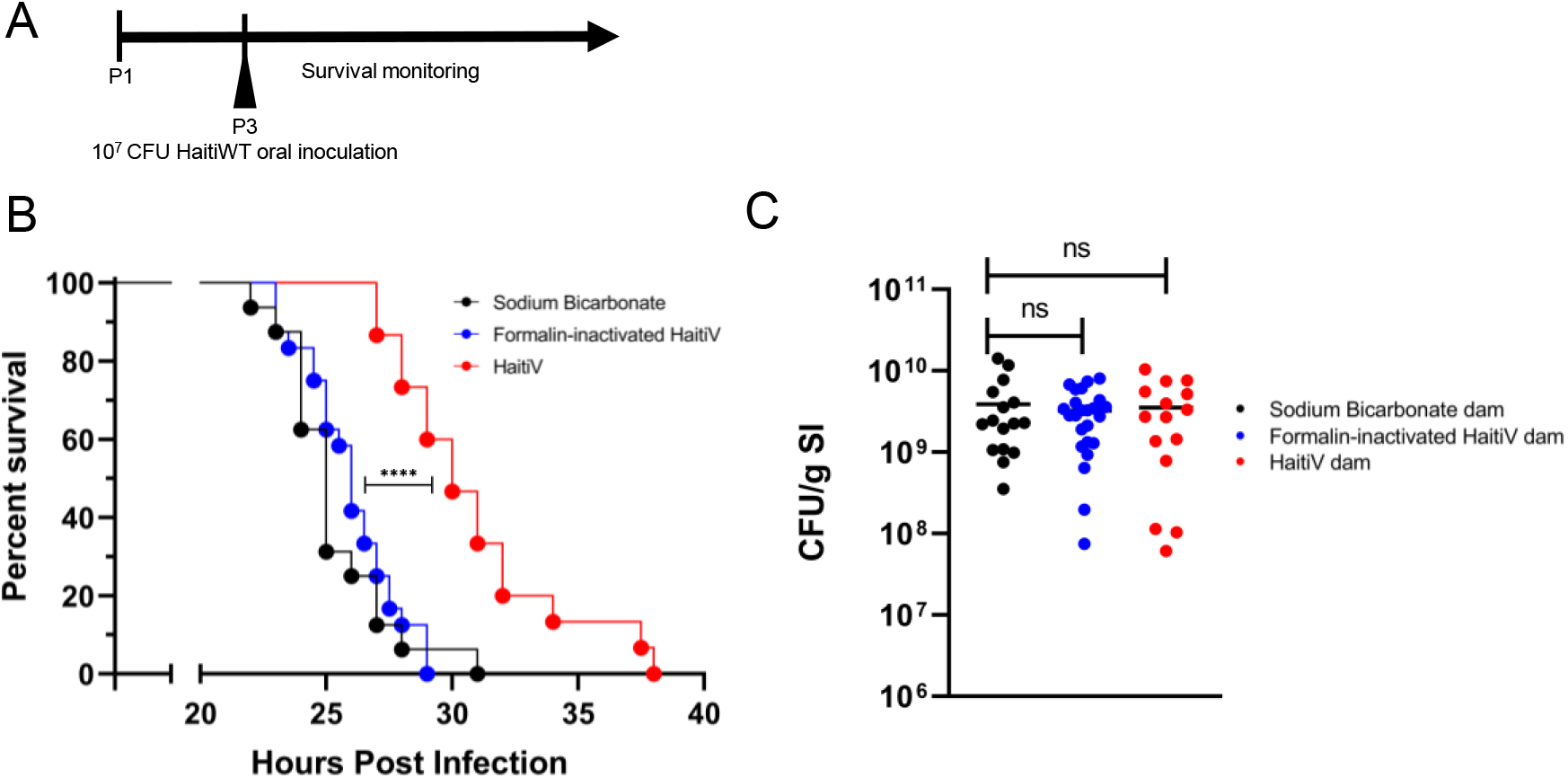
Transient colonization by HaitiV is protective in an infant mouse model of cholera. A) Survival curves of pups born to dams inoculated with HaitiV (n=15), formalin inactivated-HaitiV (n=24) or vehicle buffer (n=16) after HaitiWT challenge. Differences in the survival curves were determined by a log rank (Mantel Cox) test (p=0.0013 for HaitiV vs FI-HaitiV). B) HaitiWT CFUs recovered from the small intestines of the neonatal mice at the time they were moribund. Differences in CFU burdens were determined by a Mann Whitney U test.

Pups from HaitiV immunized dams survived significantly longer (median 30hpi) than pups from dams treated with FI-HaitiV or sodium bicarbonate control (median ~25hpi) (Fig. 4B). Despite the increased survival in the offspring of HaitiV immunized dams, at time of death, there were no differences in the burden of HaitiWT CFU recovered from the small intestines of the 3 groups (Fig. 4C). Thus, oral immunization and transient colonization of Sm-treated mice with HaitiV induces immune responses that significantly protect against choleric disease in mice.

## Discussion

The wide availability of diverse mutant mice and reagents for their study make mice a preferred model mammal for studies of human disease and therapeutics. However, adult mice are not susceptible to intestinal colonization with *V. cholerae,* confounding evaluation of OCVs. Here, we found that Sm-treated adult mice can be used to investigate the immunogenicity as well as the protective efficacy of live-attenuated OCVs. Sm-treated adult mice orally inoculated with HaitiV were colonized by this live-attenuated vaccine for 5-7days and this transient colonization was sufficient to elicit vibriocidal antibodies against both *V. cholerae* serotypes. Furthermore, pups born to HaitiV immunized Sm-treated mice exhibited prolonged survival following lethal challenge with HaitiWT compared to pups born to dams immunized with FI-HaitiV or dams treated with vehicle control. Together, these findings suggest that this model should be valuable for further studies of the efficacy and protective mechanisms of live OCVs.

Antibiotic treatment of adult mice limits the roles that the intestinal microbiota plays in inhibiting live OCV colonization. Unexpectedly, we found that oral Sm treatment enables the expansion of Sm and SXT-resistant microbes that also inhibit *V. cholerae* colonization (Fig. 3). Detailed analyses of the composition of the microbiota that are initially killed by oral Sm administration, and those that bloom after the antibiotic is withdrawn would offer valuable clues into which bacteria inhibit *V. cholerae* colonization. Though SXT treatment did not enable HaitiWT colonization of previously Sm treated animals (Fig. 3), it is possible that different antibiotic cocktails would facilitate re-challenge studies; for example, clindamycin was recently found to enable *V. cholerae* intestinal colonization of adult mice (23). Establishing conditions for re-challenge studies would simplify the model, bypassing the requirement for challenge studies in the offspring of immunized mice; however, only intestinal colonization resistance could be assayed in adult re-challenge studies since unlike suckling mice, adult mice are resistant to choleric diarrhea.

Although pups from HaitiV immunized Sm-treated dams survived longer than control animals, there was no difference in the CFU burden of HaitiWT in pups from immunized or control dams at the time they became moribund (Fig. 4). It is possible that vaccination partially delayed the replication of HaitiWT or, in addition, vaccination may elicit antibodies that antagonize toxic factors, such as cholera toxin, that promote disease, but do not directly modulate bacterial colonization. HaitiV ectopically expresses the GM1-binding B subunit of cholera toxin and immune responses to this non-toxic component of cholera toxin are linked to short term protection against cholera and enterotoxigenic *E. coli* (24).

In contrast to GF mice, which were colonized HaitiV for many months in our previous study (17),Sm-treated mice were only colonized by HaitiV for several days (Fig. 1). This is similar to the duration that live OCVs colonize the human intestine (20, 25, 26) meaning that Sm treated mice may provide a more physiologically relevant model to gauge OCV immunogenicity. However, it is important to note that although Sm-treated mice display HaitiV clearance kinetics that mimic human OCV clearance, mice are not natural hosts for *V. cholerae. V. cholerae* intestinal colonization in susceptible adult mice occurs primarily in the colon and does not rely on TCP, a critical factor for colonization of the small intestine in humans as well in infant mice and rabbits (13, 19). Nonetheless, since adaptive protective immune responses are stimulated by HaitiV in both Sm treated and GF mice, these models have value for testing vaccine immunogenicity.

We adopted the strategy that Sit et al (17) used for studying the protective efficacy of OCVs in immunized GF mice and coupled the suckling mouse model of cholera with the Sm-treated adult model of OCV immunogenicity. However, in contrast to the GF model, when the offspring of HaitiV immunized Sm-treated mice were inoculated with HaitiWT, their dams had undetectable HaitiV CFUs in their feces and VATs in their sera (Fig. 1, Fig. 4); these conditions more closely mimic those that will be present when immunized humans are exposed to *V. cholerae.* Despite the absence of circulating VATs, which are known to be relatively short-lived (27), the offspring of HaitiV-immunized dams exhibited significantly delayed death due to cholera-like disease, but all succumbed (Fig. 4). In contrast, the offspring of HaitiV-immunized GF dams were completely protected from challenge with HaitiWT (17), demonstrating the increased potency of HaitiV immunization in GF mice. The greater effectiveness of HaitiV vaccination in GF mice is likely attributable to the constant stimulation of the intestinal mucosa by HaitiV in the persistently mono-colonized GF animals and could be consistent with multiple-dose live OCV regimens. Thus, even though GF mice have immune defects, they may overestimate the potency of HaitiV and other live OCVs.

Killed whole cell vaccines like Shancol have ~50% efficacy in single dose trials (29). Notably, in marked contrast to HaitiV, we found in the Sm-treated mice model that a single dose of FI-HaitiV, which is similar to current killed whole cell vaccines, did not elicit protective immunity, consistent with the idea that live-attenuated HaitiV is far more immunogenic than killed vaccines, at least when administered as a single dose.

In summary, we have described an adult mouse model for the investigation of the protective efficacy of live OCVs. Using this model, we found that a single oral dose of HaitiV and transient colonization by the vaccine can elicit vibriocidal antibody titers and protective immune responses. Future work should enable refinement of this model to allow re-challenge of immunized animals and further characterization of these protective immune responses. Furthermore, given the availability of genetically defined mutant C57BL/6 mice, the model described here should be a valuable approach for unraveling the molecular determinants of live vaccine-mediated protection against cholera and other mucosal pathogens.

## Materials and Methods

### Bacterial Strains and culture conditions

*Vibrio cholerae* strains were grown in Luria-Bertani (LB) broth supplemented with relevant antibiotics: streptomycin (Sm) at 200μg/ml and sulfamethoxazole trimethoprim (SXT) at 80μg/mL and 16μg/mL respectively at 37°C with continuous shaking at 220rpm. For LB agar plates 1.5% agar was used and was supplemented with 5-bromo-4-chloro-3-indolyl-β-d-galactopyranoside (X-Gal) at 60μg/mL. Bacteria were kept as −80°C stocks in LB with 40% glycerol.

### Oral immunization scheme

4-week-old C57BL/6 female mice were purchased (Charles River) and housed in a BL-2 facility under conventional rearing conditions for the duration of the studies. On day 0, all mice were briefly anesthetized with isoflurane and orally gavaged with 20mg of streptomycin in 100uL of sterilized water. Mice were then provided drinking water supplemented with 5mg/mL Sm for 72 hours (22). After this, mice were then gavaged with 10^9^ CFU of an overnight culture of either HaitiV, or HaitiV inactivated by 10% formalin (FI-HaitiV) for 15 minutes, and then resuspended in 2.5% Na_2_CO_3_. Control mice were also gavaged with 100uL of 2.5% Na_2_CO_3_ alone. Mice were then provided drinking water supplemented with 200μg/ml Sm for 14 days before being returned to unsupplemented water. All mice were weighed weekly and blood samples were retrieved from each mouse by tail vein incision at each weighing. Blood samples were clotted for 45 minutes at room temperature, centrifuged at 20000xg for 10 minutes, and the serum stored at −20°C for future analysis.

Fresh fecal pellets were collected daily from each mouse, weighed, and plated in serial dilutions on LB+Sm+X-Gal agar to determine the colonization of each mouse by HaitiV. HaitiV colonizes are white due to a disrupted *lacZ*. The limit of detection of this assay represents the lowest CFU count that could be detected for a fecal pellet of that weight.

### Quantification of vibriocidal antibody titers

Circulating titers of vibriocidal antibodies were quantified by determining the minimal serum dilution required to lyse PIC158 (Ogawa) or PIC018 (Inaba) *V. cholerae* as described previously (30, 31) with minor modifications. Briefly, serial dilutions of serum were incubated with guinea-pig complement (Sigma) and the target strain, and then allowed to grow in BHI media. Reported titers are the dilution of serum that caused more than 50% reduction in target strain optical density when compared to normal saline control wells. A mouse monoclonal antibody 432A.1G8.G1.H12 targeting *V. cholerae* O1 OSP was a positive control for the assay. The limit of detection of this assay represents the lowest serum dilution at which no inhibition of growth could be detected.

### Colonization of immunized mice with toxigenic *V. cholerae*

Mice from all three groups (HaitiV, FI-HaitiV, and vehicle control) as well as 6 week old female SPF mice (Charles River) were anesthetized with isoflurane and gavaged with 16mg of sulfamethoxazole and 3.2mg trimethoprim (SXT) in 100uL of water and provided drinking water supplemented with 4mg/mL SX and 0.8mg/mL T for 72 hours. All mice were then gavaged with 10^9^ CFU of an overnight culture of HaitiWT and switched to 0.16mg/mL SX and 0.032mg/mL T in their drinking water. Fecal pellets were collected daily and plated on both SXT agar and Sm agar. Non-*V*. *cholerae* colonies were isolated by streaking on fresh Sm plates, and 16s V1/V2 DNA sequences were amplified by colony PCR using primers F341 (TCG TCG GCA GCG TCA GAT GTG TAT AAG AGA CAG CCT ACG GGN GGC WGC AG) and R805 (GTC TCG TGG GCT CGG AGA TGT GTA TAA GAG ACA GGA CTA CHV GGG TAT CTA ATC C) and sequenced. Identification of the bacteria was performed using BLAST (NCBI)

### Infant mouse survival assay

The infant mouse survival assay was adapted from a previous report (17). Female mice were mated with age matched SPF male mice (Charles River) and singly housed at E18 for delivery. At the third day of life (P3) pups were orally gavaged with 10^7^ CFU of HaitiWT and returned to their dams. Pups were monitored every 6 hours until the first signs, typically including diarrhea and dehydration, were evident. At this point, monitoring was performed every 30 minutes until pups were moribund. Pups were then removed from the nest, euthanized with isoflurane, and decapitated and dissected. The small intestines of each pup were excised, homogenized, and plated on LB agar with Sm and X-Gal.

### Statistical analysis

All statistical analyses were performed with Prism 8 (Graphpad). Infant mouse survival curves were analyzed with a log rank (Mantel Cox) test and CFU data were analyzed with a Mann Whitney U test.

### Animal use statement

All experiments in this study were performed was approved by the Brigham and Women’s Hospital IACUC (protocol 2016N000416) and in compliance with the NIH Guide for Use and Care of Laboratory animals.

## Acknowledgements

This work was supported by NIH grant (R01-AI-042347-24). BS was supported by an NSERC PGS-D fellowship (PGSD3-487259-2016). B.F., B.S., and M.K.W. designed the experiments. B.F. and B.S. performed the experiments. B.F. and M.K.W. analyzed the data. B.F., B.S, and M.K.W. wrote the manuscript. All authors reviewed and approved the manuscript.

We are grateful to Edward Ryan for providing the monoclonal antibody used for the serum assays. We thank the MKW laboratory members for helpful discussions.

M.K.W. receives funding from the HHMI and the NIH.

## Notes

### Competing Interest Statement

The authors have declared no competing interest.

