## Supplemental Figures for "Transient intestinal colonization by a live-attenuated oral cholera vaccine induces protective immune responses in streptomycin-treated mice"

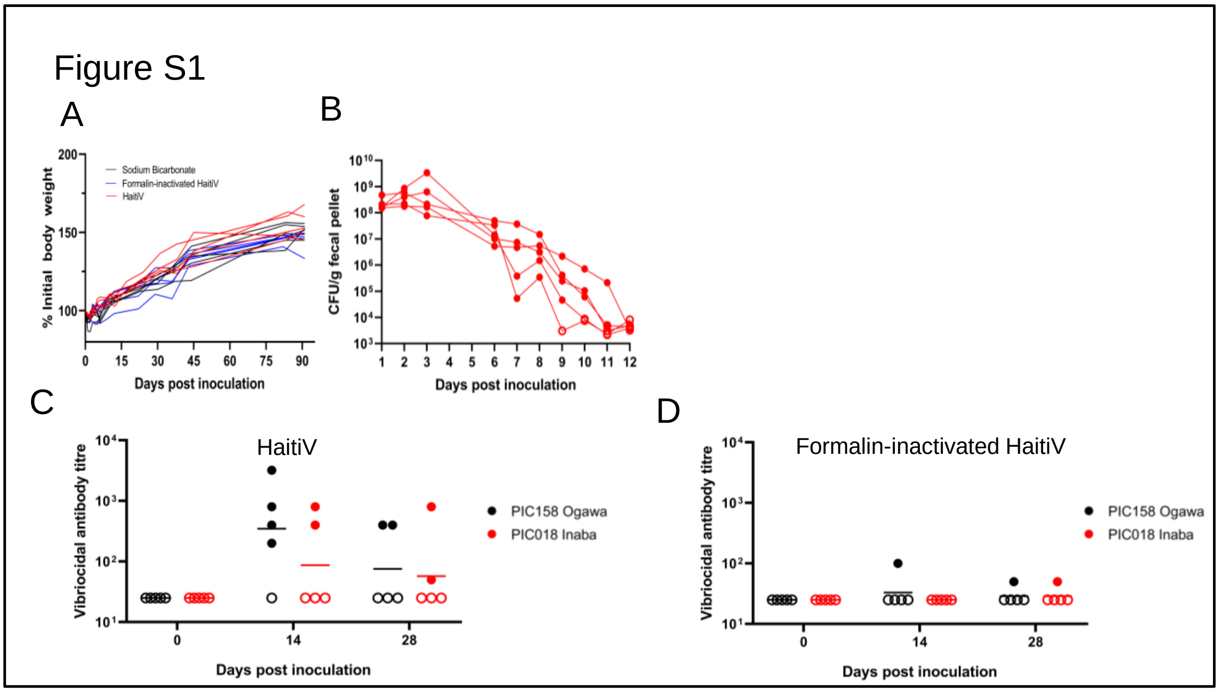


**Supplementary Figure 1. Vaccine colonization kinetics and serum vibriocidal responses in female mice that delivered pups used for the infant mouse challenges**. Mice were vaccinated according to the schematic in Fig. 1A). A) Bodyweight of all mice over the course of this study. B) HaitiV vaccinated mice fecal shedding. C) Serum vibriocidal titers from HaitiV vaccinated mice. D) Serum vibriocidal titers from formalin inactivated HaitiV vaccinated mice. Open symbols denote antibody titers below the limit of detection.
